# Stimulus-induced Gamma Power Predicts the Amplitude of the Subsequent Visual Evoked Response

**DOI:** 10.1101/388587

**Authors:** Mats W.J. van Es, Jan-Mathijs Schoffelen

## Abstract

The efficiency of neuronal information transfer in activated brain networks may affect behavioral performance. Gamma-band synchronization has been proposed to be a mechanism that facilitates neuronal processing of behaviorally relevant stimuli. In line with this, it has been shown that strong gamma-band activity in visual cortical areas leads to faster responses to a visual go cue. We investigated whether there are directly observable consequences of trial-by-trial fluctuations in non-invasively observed gamma-band activity on the neuronal response. Specifically, we hypothesizedthat the amplitude of the visual evoked response to a go cue can be predicted by gamma power in the visual system, in the window preceding the evoked response. Thirty-three human subjects (22 female) performed a visual speeded response task while their magnetoencephalogram (MEG) was recorded. The participants had to respond to a pattern reversal of a concentric moving grating. We estimated single trial stimulus-induced visual cortical gamma power, and correlated this with the estimated single trial amplitude of the most prominent event-related field (ERF) peak within the first 100 ms after the pattern reversal. In parieto-occipital cortical areas, the amplitude of the ERF correlated positively with gamma power, and correlated negatively with reaction times. No effects were observed for the alpha and beta frequency bands, despite clear stimulus onset induced modulation at those frequencies. These results support a mechanistic model, in which gamma-band synchronization enhances the neuronal gain to relevant visual input, thus leading to more efficient downstream processing and to faster responses.

**Significance statement:** Gamma-band activity has been associated with many cognitive functions and improved behavioral performance. For example, high amplitude gamma-band activity in visual cortical areas before a go cue leads to faster reaction times. However, it remains unclear through which neural mechanism(s) gamma-band activity eventually affects behavior. We tested whether the strength of induced gamma-band activity affects evoked activity elicited by a subsequent visual stimulus. We found enhanced amplitudes of early visual evoked activity, and faster responses with higher gamma power. This suggests that gamma-band activity affects the neuronal gain to new sensory input, and thus these results bridge the gap between gamma power and behavior, and support the putative role of gamma-band activity in the efficiency of cortical processing.

## Introduction

Mesoscopic and macroscopic electrophysiological signals, as measured invasively as local field potentials (LFPs) or non-invasively as the magneto/electroencephalogram (MEG/EEG), can often be characterized by rhythmic activity patterns in a broad range of frequencies (Buzsáki and Draguhn, 2004). Experimentally, distinct frequency bands have been implicated in various cognitive processes. For instance, cortical gamma-band activity (30-90 Hz) has been associated with attention (Tiitinen et al., 1993; Fries et al., 2001; Taylor et al., 2005), memory (Jensen and Lisman, 1996; Carr et al., 2012) and perception (Gray and Singer, 1989; Llinas et al., 1994). Gamma rhythms result from a balanced interplay between neuronal excitation and inhibition. Fast-spiking interneurons bring about the inhibition of the excitatory drive within a population. Once the inhibition fades off, the excitatory drive activates pyramidal cells and in turn, excites the feedback loop of fast-spiking interneurons. This interaction synchronizes the IPSPs in pyramidal neurons and generates gamma rhythms at the population level (Buzsáki and Wang, 2012). Fries (2015) proposed that this mechanism functions to synchronize inputs down the processing hierarchy, thereby making communication between neuronal groups more effective.

Given its putative mechanistic role in affecting the outcome of cortical computations, gamma-band synchronization has become a popular neural substrate to quantify in relation to behavior during cognitive experiments. This has led to evidence for a relationship between gamma-band synchronization and behavior, both in humans and other mammals. Multiple studies have found a larger pre-stimulus gamma power for perceived versus unperceived stimuli (Makeig and Jung, 1996; Linkenkaer-Hansen, 2004; Hanslmayr et al., 2007; Wyart and Tallon-Baudry, 2008), and strong gamma power in visual areas leads to faster responses to a visual go cue (Womelsdorf et al., 2006; Koch et al., 2009; Hoogenboom et al., 2010). These results are in line with the idea that gamma-band synchronization facilitates stimulus processing, and more specifically, they suggest a behavioral relevance of the strength of gamma-band synchronization in task-relevant areas.

Although the relation between the amplitude of the gamma rhythm and behavior has been established, relatively little is known about how gamma-band synchronization in sensory cortical areas affects the chain of neuronal events, leading to an eventual effect on behavior. One way to investigate this would be to relate trial-by-trial fluctuations of the gamma amplitude and/or phase, estimated at the moment of task-relevant stimulus onset with the transient event-related response to this stimulus. Most studies investigating the mechanisms of gamma-band facilitation used invasive recording techniques, and focused on the relevance of the gamma phase (Fries et al., 2001; Cardin et al., 2009; Ni et al., 2016). Ni et al (2016) show, at the mesoscopic scale of LFPs and multiunit activity, that gamma-band oscillations lead to rhythmic fluctuations in neuronal gain, such that inputs at phases of high gain elicit stronger multiunit activity.

In the present research, we investigated the effect of trial-by-trial fluctuations in MEG-derived gamma-band activity on stimulus-evoked activity. Using a visual stimulation paradigm that is known to robustly induce gamma-band activity in early visual cortical areas, we instructed participants to respond as fast as possible to an unpredictable salient change in a moving grating. We hypothesized that intrinsic variability in gamma power reflects variability in the efficiency of information transfer in the visual processing stream, which would manifest itself as correlated amplitude variability of the early evoked responses. More salient activation in sensory areas would in turn lead to enhanced processing in downstream areas, eventually causing a faster behavioral response.

## Materials and Methods

### Subjects

33 healthy volunteers, of which 22 females and 11 males, participated in the study. Their age range was 18-63 years (mean ± SD: 27 ± 10 years) and they all had normal or corrected-to-normal vision. All subjects gave written informed consent according to the Declaration of Helsinki. The study was approved by the local ethics committee (CMO region Arnhem/Nijmegen). One subject was excluded from analysis due to a technical error, which corrupted one of the data files.

### Experimental Design

#### Stimuli

The experimental task was programmed in MATLAB (R2012b, Mathworks, RRID: SCR_001622) using Psychophysics Toolbox (Brainard and Vision, 1997, RRID: SCR_002881). All stimuli were presented against a gray background. A fixation dot was present throughout the experiment, the color of which indicated when the participant was allowed to blink with their eyes (green for blinking, red for not blinking). A concentric sinusoidal grating was presented at 100% black/white contrast and was tapered towards the edges with a Hanning mask, such that edge effects were excluded (see figure 1). The grating was present at the center of the screen, with a visual angle of 7.1°, 2 sinusoidal cycles per degree and a contraction speed of 2 cycles per second.

**Figure 1.**
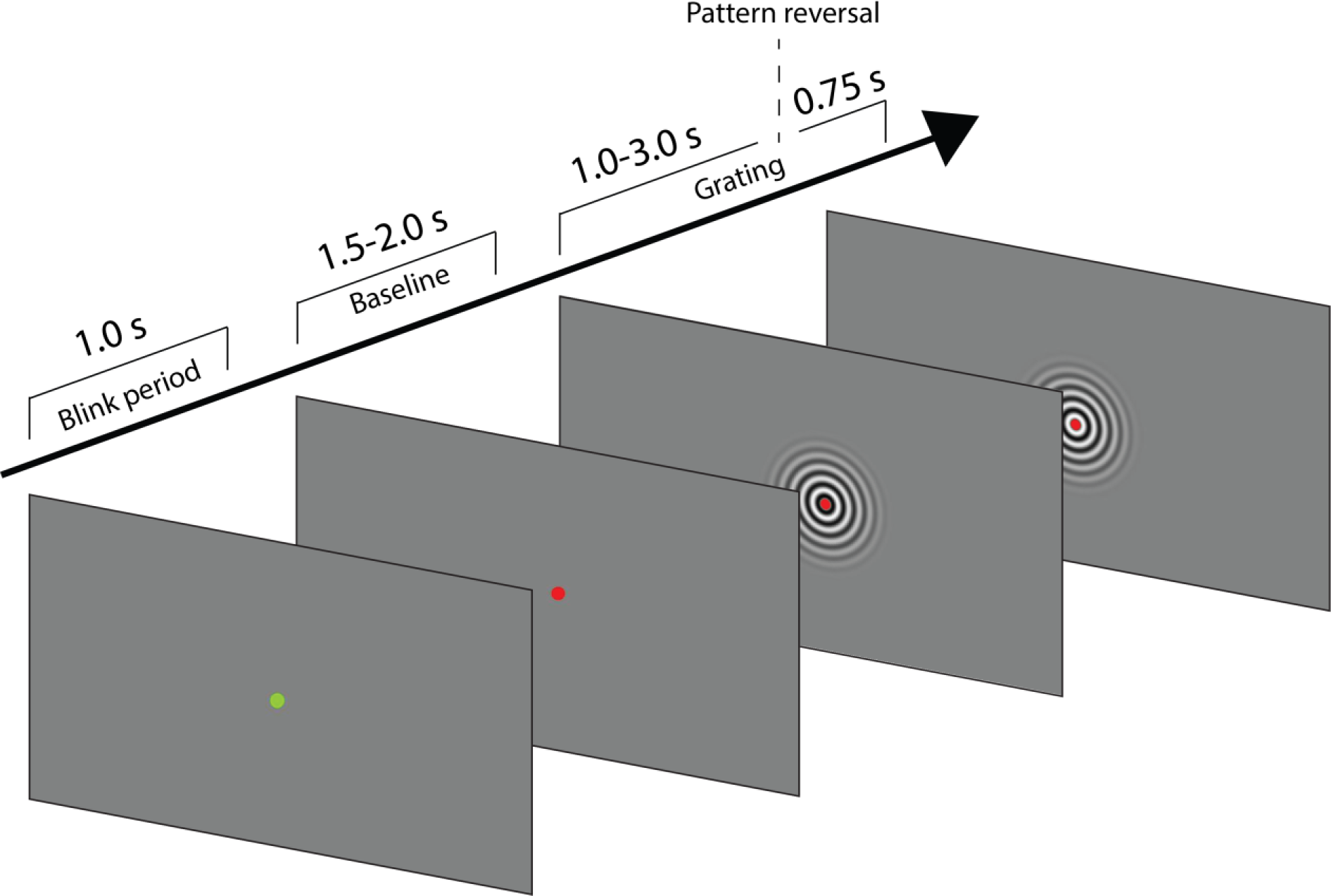
Task time-line. Each trials starts with a 1.0 second blink period, followed by a variable baseline period (1.5-2.0 s). A concentric drifting grating is presented for 1.0-3.0 seconds, after which a stimulus reversal occurs. The grating continues drifting for 0.75 seconds, until the end of the trial, during which a response has to be made.

#### Experimental equipment

Stimuli were presented by back-projection onto a semi translucent screen (width 48 cm) by an PROPixx projector with a refresh rate of 120 Hz and a resolution of 1920 x 1080 pixels. Subjects were seated at a distance of 76 cm from the projection screen in a magnetically shielded room. MEG was recorded throughout the experiment with a 275-channel axial gradiometer CTF MEG system at a sampling rate of 1200 Hz. In addition, subject’s gaze position was continuously recorded using an SR Research Eyelink 1000 eye-tracking device (RRID: SCR_009602). Head position was monitored in real-time during the experiment by using head-positioning coils at the nasion and left and right ear canals of the subject (Stolk et al., 2013). When head position deviated more than 5 mm from the position at the start of the experiment, subjects readjusted to the original position. Behavioral responses during the MEG session were recorded using a fiber optic response pad (FORP).

In addition to the MEG recording, anatomical T1 scans of the brain were acquired with a 3T Siemens MRI system (Siemens, Erlangen, Germany). In order for co-registration of the MEG and MRI datasets, the scalp surface was mapped using a Polhemus 3D electromagnetic tracking device (Polhemus, Colchester, Vermont, USA).

#### Procedure

Subjects were instructed to keep fixation at the fixation dot throughout the experiment (see figure 1). The fixation dot was colored red most of the time, but turned green during the eye-blink period. After a 1.0 second eye-blink period and a 1.5-2.0 second baseline window, a contracting grating was presented at the center of the screen. The grating contracted for 1.0-3.0 seconds, after which a pattern reversal of the stimulus occurred. This functioned as a go cue. Participants had to respond as fast and as accurately as possible to the go cue by pressing a button with the right index finger. Responses had to be made within 700 ms. Ten percent of the trials were catch trials, in which no stimulus change occurred. After the stimulus change the grating continued to contract for another 750 ms, until the end of the trial. There was no feedback of task performance, but participants were trained before the experiment to make sure they understood the task. Participants completed a maximum of thirteen blocks, each consisting of forty trials, or until one hour had passed. In between blocks there was a self-paced break, if needed followed by repositioning of the subject to the original position (see Experimental Equipment). In total, participants completed between 400 and 520 trials.

### Data Analysis

#### MEG preprocessing

The MEG data was preprocessed offline in MATLAB (2015b, Mathworks, RRID: SCR_001622) using FieldTrip toolbox (Oostenveld et al., 2011, RRID:SCR_004849) and custom written code. First, excessively noisy channels and trials were removed from the data by visual inspection. Additionally, trials with squid jumps or muscle artifacts were removed from the data. Eyetracker data was visually inspected to discard trials with eye blinks within the latency of interest and trials where the eye position exceeded 5 degrees from the fixation dot were removed likewise.

The data were demeaned, and high pass filtered at 1 Hz using a finite impulse response windowed sinc (FIRWS; Widmann, 2006) filter. Power line interference (50 Hz) and its harmonics were removed using a discrete Fourier transform (DFT) filter. Further, signals relating to cardiac activity or eye blinks and eye movements were identified and removed from the data using independent component analysis (ICA). Lastly, the trials of interest were defined as those where a stimulus change was present and where a behavioral response was made within 700 ms of that event.

#### MRI processing

MRI data were co-registered to the MEG-based coordinate system using the head-positioning coils and the digitized scalp surface. Using SPM8 (Penny et al., 2011) we created volume conduction models of the head, and individual meshes of dipole positions, consisting of a cortically constrained surface-based mesh with 15,784 vertex locations. These meshes were created using Freesurfer (RRID: SCR_001847) and HCP workbench (RRID: SCR_008750). The dipole positions were used for the identification of a virtual channel with the strongest gamma-band response or low frequency response (see *Single-trial power*), and evoked responses were modeled on a parcellated version of this mesh. The vertices were grouped into 374 parcels based on a refined version of the Conte69 atlas (Van Essen et al., 2012) in order to reduce the dimensionality of the data (similar to Schoffelen et al., 2017). Forward models were computed using single-shell volume conductor models that were derived from individual structural MR images (Nolte, 2003).

#### Time-frequency analysis

Time-resolved spectral power was estimated for low (2-30 Hz) and high (28-100 Hz) frequencies after padding the data with zeros to six seconds. For low frequencies, a Hanning tapered 500 ms sliding time window was used in steps of 50 ms, with 2 Hz resolution. High frequency power was estimated using a DPSS multi-taper approach with a sliding time window of 250 ms and steps of 50 ms, 4 Hz resolution and 8 Hz smoothing. Time-frequency activity was expressed relative to a baseline, defined as [-1.0 −0.25] seconds, time locked to stimulus onset. For initial exploration, spectral decomposition was performed on synthetic planar gradient data (Bastiaansen and Knösche, 2000), and combined into a single spectrum per sensor. This way, power spectra could easily be averaged across subjects for visualization purposes.

#### Peak frequency

Subject-specific gamma power was estimated on the individualized gamma peak frequency. In order to estimate peak frequencies, the power spectrum after stimulus onset was contrasted with the pre-stimulus baseline. First, trials were separated into baseline and stimulus presentation epochs, where the first 400 ms of stimulus presentation were discarded in order to prevent spectral effects of evoked activity. Next, trial epochs were cut into 500 ms snippets, with fifty percent overlap. Spectral power was then estimated on these snippets in the 30-90 Hz range, after tapering with a Hanning window. Finally, the gamma peak frequency was determined at the maximum power ratio of stimulus presentation over baseline period, averaged over occipital MEG channels. A similar approach was used for low frequencies, in the 2-30 Hz range, but here the negative peak (i.e. showing the largest power reduction from baseline) was used.

#### Single-trial power

To get an optimal estimate of single-trial gamma power, we created subject-specific virtual channels, using a DICS beamformer (Gross et al., 2001), scanning the cortically constrained mesh of dipole positions. The 750 ms of data before the stimulus change were used to ensure the best signal-to-noise ratio, while at the same time ensuring that the estimate of gamma-band activity was as little as possible affected by evoked activity. These 750 ms epochs were padded with zeros to one second, and a multi-taper Fast Fourier transform (FFT) with 8 Hz spectral smoothing was applied to these data. The same was done for a 750 ms baseline window. Spatial filters were created for each of the vertex locations, using the cross-spectral density estimated from the concatenated data, at the subject-specific peak frequencies, and a regularization parameter of 5%. Next, the virtual channel was selected as the vertex that showed the largest increase in gamma power, relative to baseline.

Next, the single-trial gamma power was estimated on these virtual channels, in the 200 ms just before the ERF window (see *Single-trial event-related responses*; the window in which power was estimated ended 20 ms before the start of the ERF window), using a spatial filter with fixed dipole orientation, optimized for this time window.

To estimate single-trial power estimates for the alpha-beta band, a similar procedure was used. The cross-spectral density matrix was estimated at individual peak-frequency with 2.5 Hz smoothing, based on the 400 ms before the ERF window. Since the induced low frequency response was a power decrease relative to baseline, the vertex location that showed the largest decrease was selected as the virtual channel.

#### Single-trial event-related responses

The event-related response to the stimulus change was modelled using a Linearly Constrained Minimum Variance (LCMV) beamformer on the cortically-constrained meshes of dipole positions, followed by a parcellation based on an anatomical atlas (see *MRI preprocessing*). Data, time locked to stimulus change, were selected and baseline corrected based on 100 ms prior to the change. For each anatomical parcel, the source time courses of the dipoles belonging to this parcel were concatenated, and subjected to a principal component analysis (PCA). The first spatial component that explained most variance in the signal was used as a representation of single-trial activity for this parcel. In order to account for the beamformer’s depth bias, the data were normalized by an estimate of the noise using the covariance matrix of the 200 ms prior to the go cue. The resulting time courses were low-pass filtered at 30 Hz using a finite impulse response (FIR) windowed sync function. The data were filtered from right to left, to avoid leakage of pre-change signal into the post-change estimates. Next, the amplitudes of the single-trial visual evoked responses were estimated in a time window showing the most prominent peak in the trial averaged ERF, in the first 100ms after stimulus change, on a subjects-by-subject basis. These windows were manually defined by visual inspection of the source-level activity time courses.

#### Correlation of single-trial power and ERF amplitude

Correlations between ERF amplitude, alpha-beta power and gamma power, and response speed, and between gamma power, response speed and trial length were computed at the single-subject level using Spearman’s rank correlation coefficient. We also computed partial correlations between alpha-beta/gamma power and response speed, each time accounting for trial length and power values in the other frequency band.

#### Statistical analysis

The correlation between alpha-beta power and gamma power, response speed and trial length was statistically evaluated using a parametric t-test (against 0) on the distribution of correlation coefficients over subjects (alpha = 0.05). Statistical significance of the correlation between alpha-beta/gamma power and ERF amplitude, and between ERF amplitude and reaction times was assessed using non-parametric permutation tests (based on 10000 permutations) combined with spatial clustering for family-wise error control (Maris and Oostenveld, 2007). Under the null hypothesis of no systematic relationship across participants between gamma power and the ERF amplitude, we created a reference distribution of the group-level t-statistic of the correlation against zero, using sign swapping of the correlation for random subsets of subjects. Spatially adjacent parcels with t-values corresponding to a nominal alpha threshold of 0.05 (0.01 for the correlation between ERF amplitude and RT) were grouped into clusters, and cluster-level statistics were computed as the sum of t-values within a cluster. The null-hypothesis was rejected if the maximum cluster-level statistic in the observed data was in the positive tail of the permutation distribution of cluster-level statistics for the correlation between ERF amplitude and gamma power, and in the negative tail for the correlation between alpha-beta power and ERF amplitude, and between ERF amplitude and reaction times, at a level of <0.05 one-sided.

## Results

Out of the 400-520 completed trials per subject, on average fifty of them were catch trials, i.e. these trials did not require a response. Overall, the subjects performed with a mean accuracy of 94% (SD = 5.8%). Excluding catch trials and trials with artifacts or excessive eye movements, on average 339 trials per subject (SD = 61) were considered for further analysis. Of this pool, the mean performance rate was 94% (SD = 5.5%) and the mean reaction time over subjects was 371 ms (SD = 56 ms).

### Stimulus protocol leads to reliable stimulus-induced changes in gamma power and visual event-related responses

The contracting grating stimuli used here are known to robustly induce gamma-band synchronization (Hoogenboom et al., 2006; Swettenham et al., 2009; Van Pelt and Fries, 2013). In order to verify the spectral characteristics of the stimulus-onset induced neuronal response, we conducted a time-frequency analysis at the channel-level. We contrasted spectral power after stimulus onset with the average spectrum in the baseline window. Figure 2 shows the average spectral power for all subjects in the gamma-frequency range (30-90 Hz). As expected, it was highest in occipital channels and it remained high throughout the whole stimulus presentation. Sources of the activity were localized to early visual areas (see figure 2e). In order to assess the gamma power increase quantitatively we estimated the power increase from baseline at the individual gamma peak frequency (figure 2d) and at the occipital channel that showed maximal increase. Over subjects, gamma power increased on average with 124% (mean, SD =124%) during stimulation (figure 2d, right). Besides a gamma-band increase, a decrease in power was generally observed in the alpha/low beta band (8-20 Hz). This phenomenon presented itself mainly in occipital channels, and was also strongest in occipital source parcels (see figure 3).

**Figure 2.**
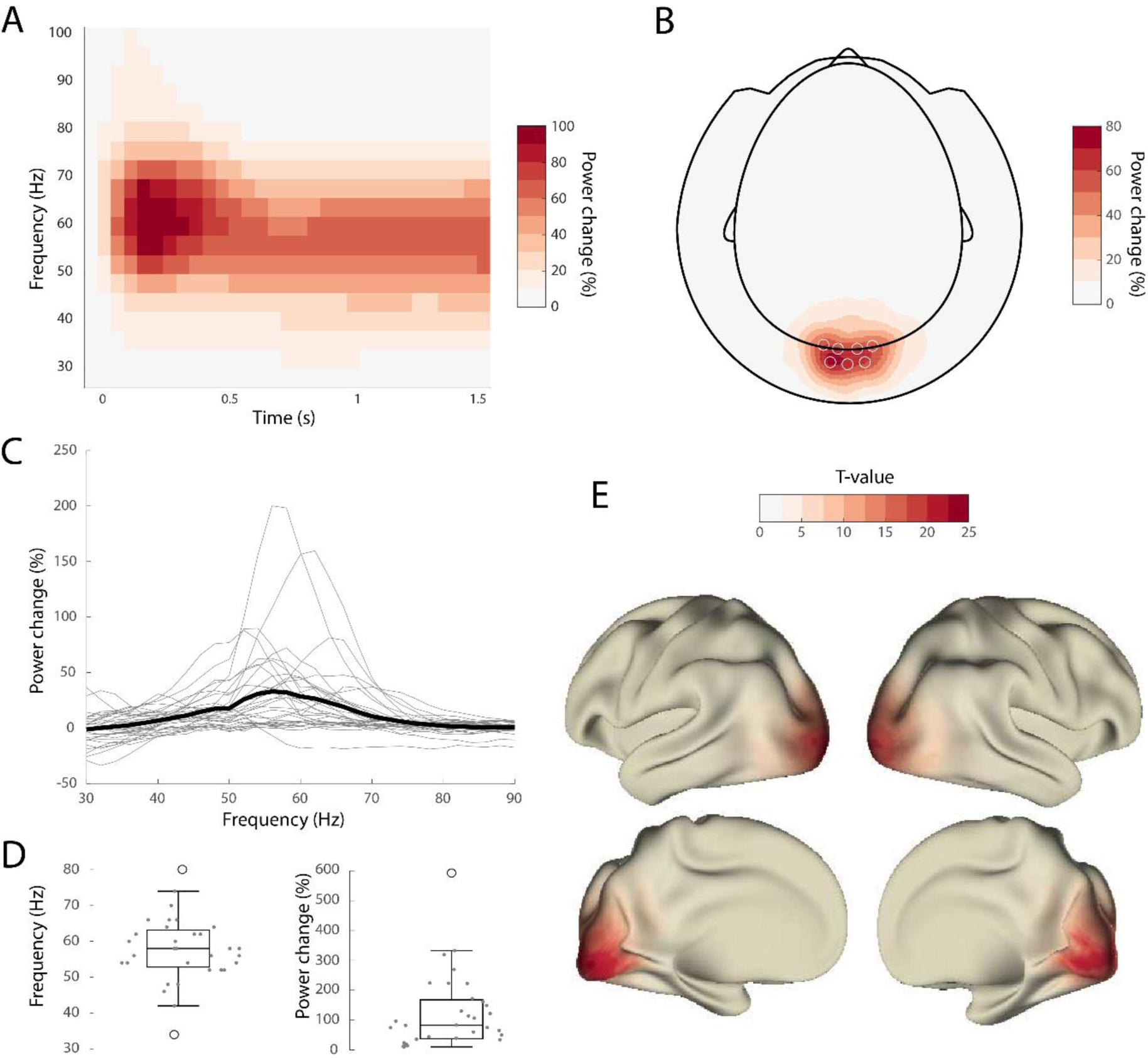
Stimulus induces strong gamma synchronization. A) Group-average time-frequency spectrum. Gamma (50-70 Hz) power peaks right after stimulus onset and is sustained throughout stimulus presentation. B) Topography of the stimulus-induced gamma-band activity. Circles reflect selected channels, shown in A. C) Individual (gray line) and group-average (black line) power spectra on the channel level. D) Left: gamma peak frequencies. Generally, peak frequencies were in the 40-70 Hz range. Right: power changes (right) at individual gamma peak frequencies. Power changes were highly variable. E) Source level activity of induced gamma the group average t-value. Induced gamma activity was strongest in occipital regions.

In addition to the stimulus inducing a robust gamma-band response, the stimulus change caused an event-related field (ERF). Figure 3a shows the ERF of an example subject over trials, together with the topography of the trial average of the P100 response in figure 3c. The signal-to-noise ratio (SNR) of the evoked response is relatively poor at the single-trial level. Also, the spatial topography was variable over subjects (data not shown). In order to reduce the spatial variability over subjects and to boost the SNR, all further analyses were conducted on the source level. Figure 3b shows a superior SNR for single trials on the source level compared to the channel level, together with the source topography in figure 3d. Despite the large variability of the response evoked by the stimulus change and the percentage increase in induced gamma, all subjects did show the neuronal response that was expected.

**Figure 3.**
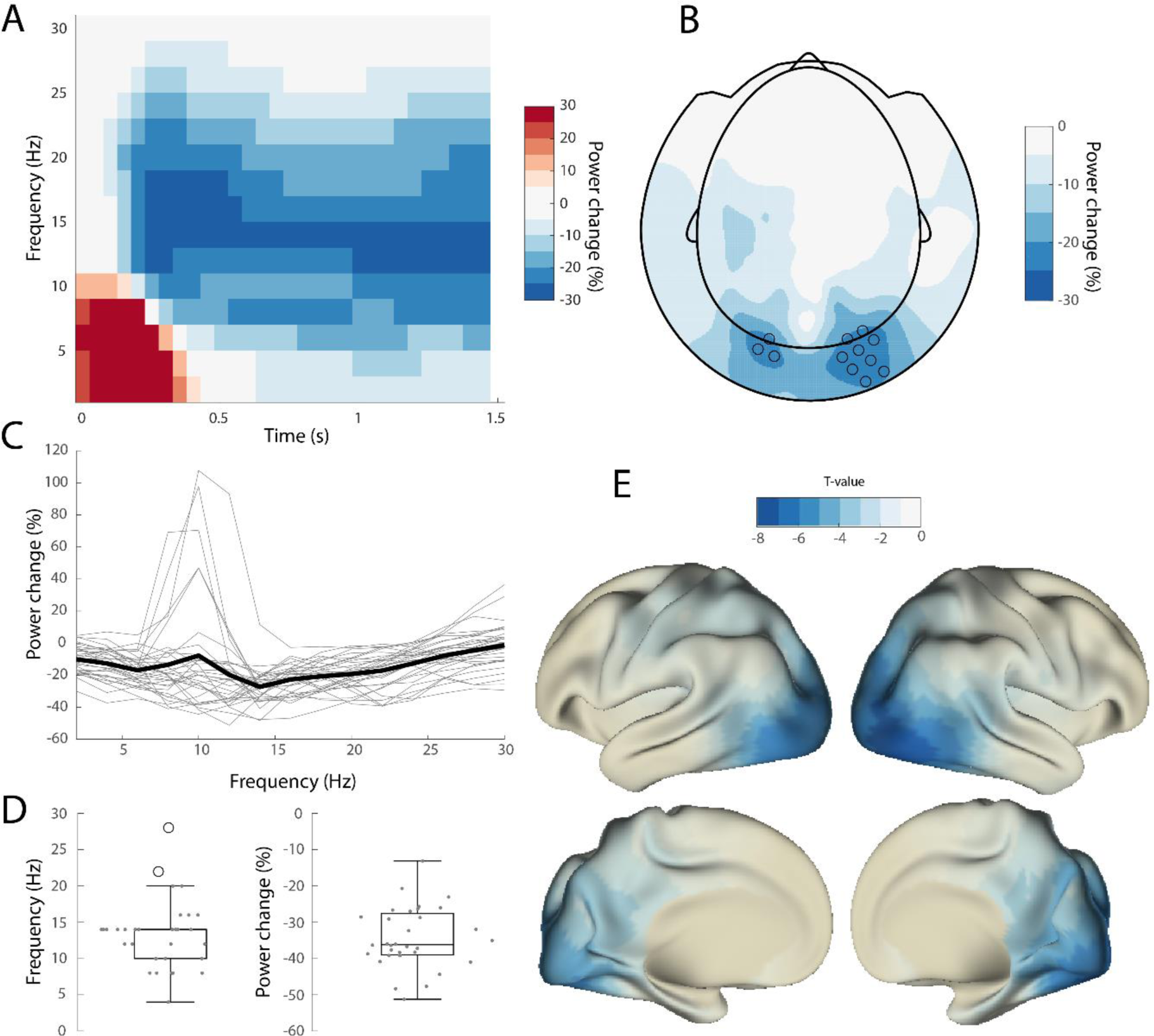
Stimulus induces desynchronization in alpha-beta band (8-20 Hz). Similar to figure 2 but for low frequencies. Power in de alpha and low beta range decreased after stimulus onset, which was 286 strongest in occipital parts of the cortex.

**Figure 4.**
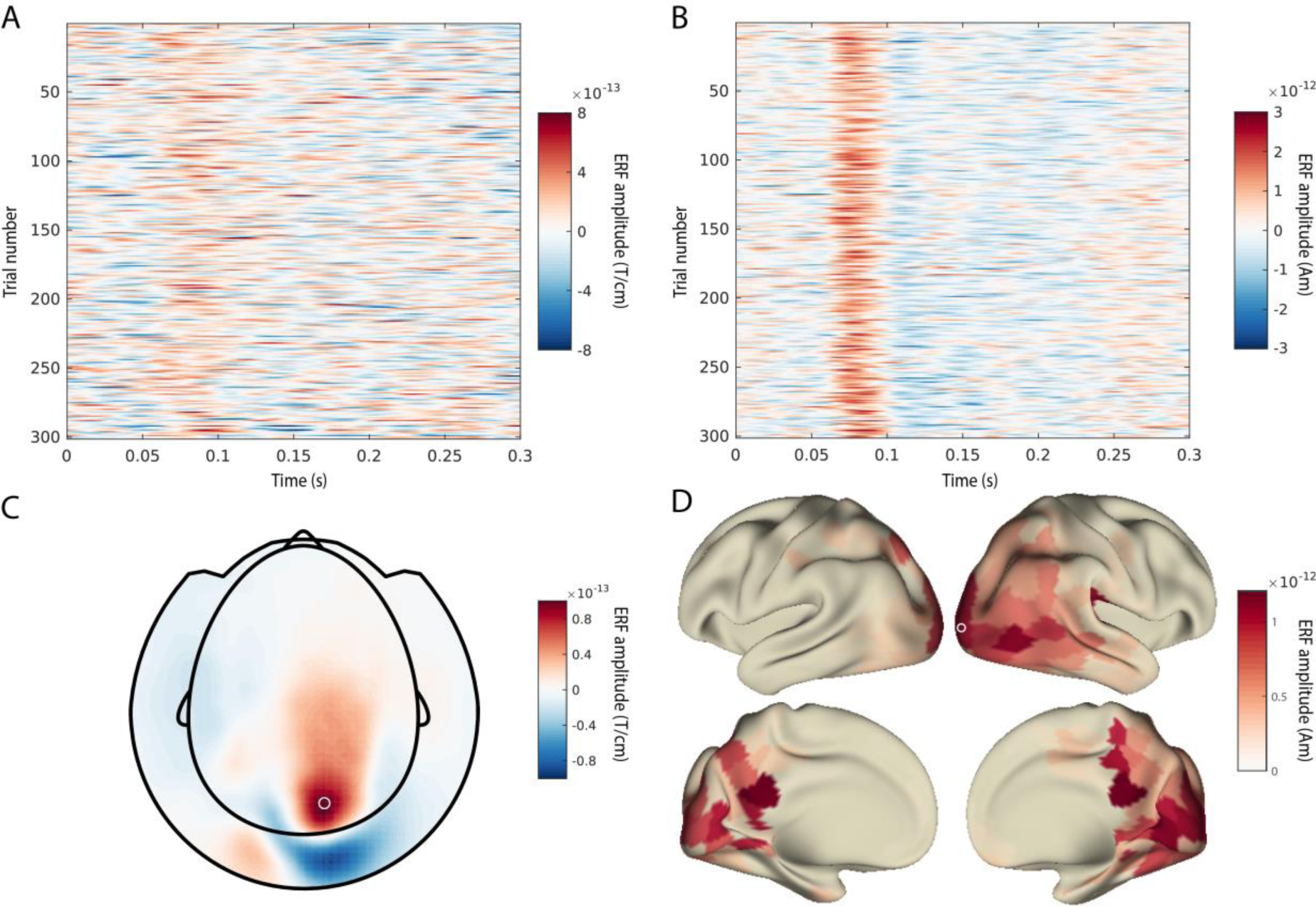
Signal-to-noise ratio of evoked activity is higher at the source level than at the channel level. A-B: single-trial time courses of evoked activity for the channel/parcel depicted in C/D, for a representative subject. C-D: topography of the P100 (depicted in A/B), evoked by the stimulus change.

### Pre-stimulus gamma power correlates with reaction times

In order to evaluate the effect of pre-stimulus gamma power on behavior, we calculated the correlation between reaction times and the gamma power preceding the stimulus change. Gamma power was estimated on a virtual channel in source space. There was a relatively weak but highly significant, negative correlation of −0.072 (SD = 0.064, t(31) = −6.3, p = 5.1*10^−7^), which is in line with previous research (Womelsdorf et al., 2006; Hoogenboom et al., 2010).

One potential factor that might explain the correlation between gamma power and reaction times is stimulus jitter (i.e., the time between stimulus onset and the stimulus change). Stimulus jitter was uniformly distributed, and by consequence the instantaneous probability of a reversal event (hazard rate) increased over time. There could be a common dependence of gamma power and reaction times on stimulus expectancy. In order to investigate this possibility, we computed partial correlations between reaction time, gamma power and stimulus jitter, each time accounting for the third variable. Since we also found a stimulus-induced power reduction in the alpha-beta band, there is also a possibility that the correlation between gamma power and reaction times is actually caused by a common dependence on power in this frequency band. Therefore, we also partialled out power in the alpha-beta band. Reaction times correlated negatively with both gamma power (M = −0.068, t(31) = −6.1, p = 8.9 * 10^−7^, uncorrected) and stimulus jitter (M = −0.17, t(31) = −7.0, p = 6.9 * 10^−8^, uncorrected), but there was no correlation between gamma power and stimulus jitter (t(31) = 0.85, p = 0.40, uncorrected). Additionally, no significant correlation was found between reaction times and low frequency power when accounting for gamma power and stimulus jitter (t(31) = 1.1, p= 0.27, uncorrected), nor between low frequency power and gamma power (t(31) = 0.03, p = 0.97, uncorrected). These results indicate that the correlation between gamma power and reaction times is not likely to be the result of a build-up in expectancy, nor a result of power correlations between frequency bands. Further, there is no effect of low frequency power on reaction times, above and beyond the effect of gamma.

### Pre-stimulus gamma power predicts ERF amplitude

Considering the vast amount of variability in the evoked response over subjects at the channel level, and the low signal-to-noise ratio of single trial event-related responses, we estimated the single trial responses to the stimulus change at the source-level, before quantifying the relation between gamma power and ERF amplitude. The time courses of the evoked response were projected into source space with an LCMV beamformer and combined into parcels according to an anatomical brain atlas (Van Essen et al., 2012). The relation between gamma power and the ERF was quantified as a Spearman rank correlation at the single subject level and can be seen in figure 5. Group statistical evaluation showed that the correlation differed significantly from zero (M = 0.027, p=0.01, nonparametric permutation test, corrected). This difference was supported by a cluster of positive correlation in source parcels in occipital and parietal areas (figure 5a), supporting the hypothesis that increased gamma power leads to an increased amplitude of the stimulus-evoked transient.

**Figure 5.**
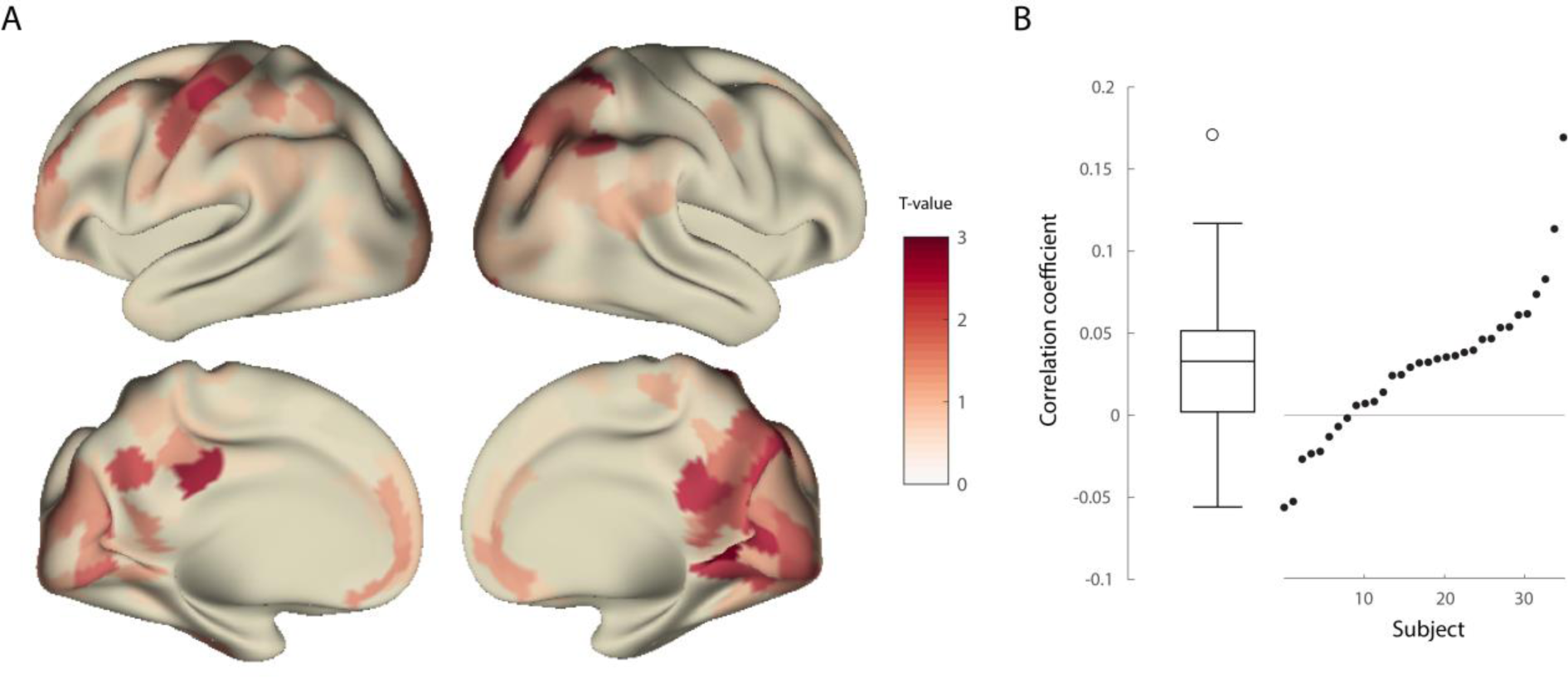
Occipital gamma power correlates with ERF amplitude. A) Group statistic of the correlation between gamma power and ERF amplitude. B) boxplot (left) of correlation values and individual correlations (right). For each subject, correlations were averaged over those parcels belonging to the cluster that contributed to the significant effect.

Additionally, we correlated ERF amplitude and reaction times (figure 6b). This correlation was significantly lower than zero (M = −0.027, p=0.037, nonparametric permutation test, corrected), indicating that a higher amplitude of early evoked activity leads to a faster behavioral response. The cluster that mostly contributed to this effect was found in source parcels belonging to the visual cortex of the right hemisphere (figure 6a), conform our hypothesis. Although we did not find a correlation between low frequency power and reaction times, low frequency power might still affect ERF amplitude. In order to ensure that this was not the case, we tested whether low frequency power before the ERF was predictive of ERF amplitude. Conform the stimulus-induced alpha-beta power decrease, if any, a negative correlation was expected with ERF amplitude. This correlation was not significant at the group level (p = 0.66, nonparametric permutation test, corrected).

**Figure 6.**
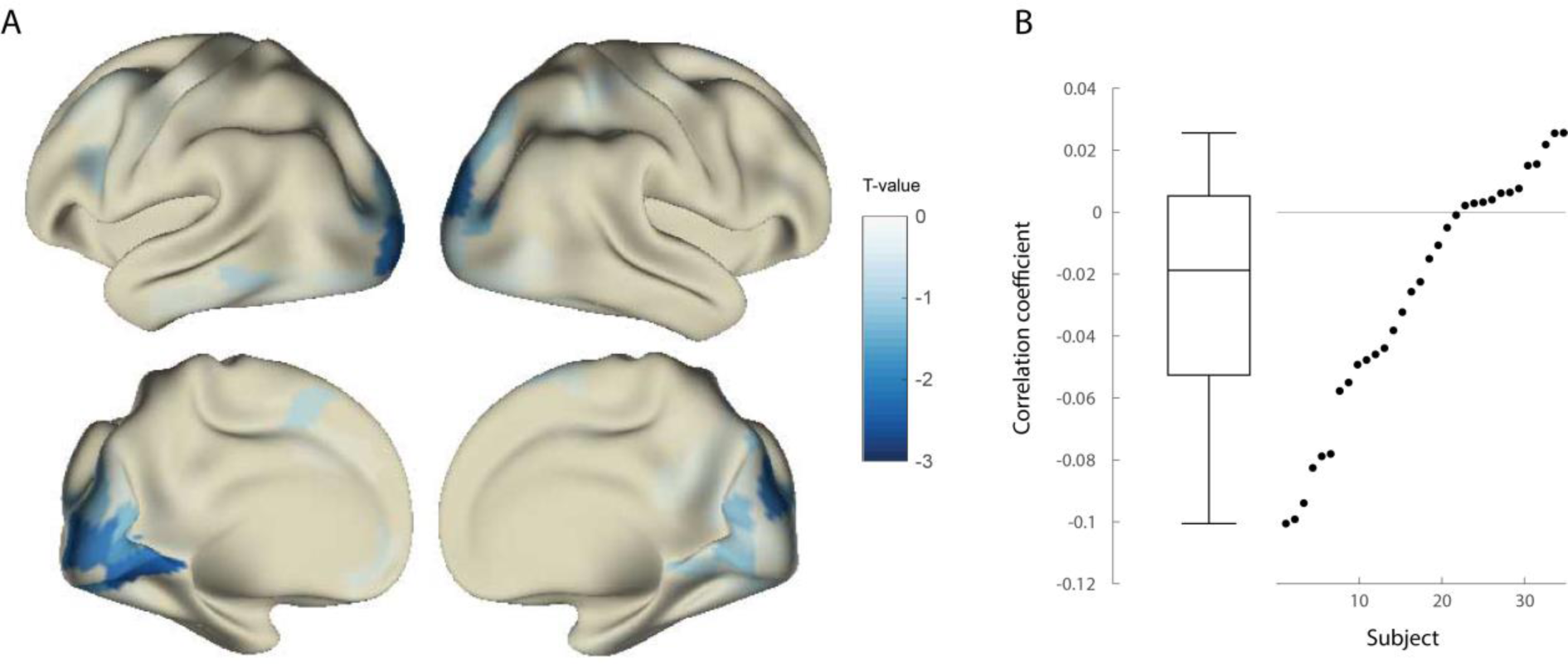
ERF amplitude in occipital cortex correlates with reaction times. A) Group statistic of the correlation between ERF amplitude and reaction times. B) boxplot (left) of correlation values and individual correlations (right). For each subject, correlations were averaged over those parcels belonging to the cluster that contributed to the significant effect.

## Discussion

In this experiment, we investigated the neuronal consequences of trial-by-trial variability in induced gamma-band activity, using a visual stimulus change detection paradigm. We hypothesized that higher gamma power before a go cue would facilitate the efficiency of the processing of the response cue, leading to a more strongly synchronized response in early visual areas, as reflected by higher early latency ERF amplitudes. In turn, the increased processing efficiency would lead to a faster behavioral response.

We computed single trial estimates of gamma power in visual areas, in the time window just prior to the stimulus change, and replicated the finding that higher gamma amplitude leads to faster reaction times in response to the go cue (Hoogenboom et al., 2010). Moreover, we correlated the single-trial gamma power with the amplitude of early latency source-reconstructed event-related activity, and observed a significant group-level effect, where the early latency event-related response in parieto-occipital areas correlated positively with pre-stimulus gamma power. In turn, the amplitude of event-related response in visual areas correlated negatively with reaction times. These findings are consistent with the hypothesis that strong local gamma-band synchronization facilitates the neuronal response to a change in the stimulus, which eventually leads to improved behavioral performance.

Here, the effect of oscillatory activity on the subsequent neuronal response was specific to the gamma band. We also analyzed the effect of alpha/beta activity on the event-related response, since activity in these frequency bands was also prominently modulated by the onset of the visual stimulus. In contrast to the gamma band, we did not observe a significant association between trial-by-trial power fluctuations in these lower frequencies, and trial-by-trial fluctuations in the event-related response, and response speed.

Although Hoogenboom et al. (2010) did not specifically investigate the relation between gamma-band activity and the ERF, the authors did quantify the relation between ERF amplitude and reaction times, and found no significant effect. This latter null-finding is in contrast with our analysis of the current data. Most likely this discrepancy is due to the fact that the aforementioned study used a temporally ill-defined stimulus change (change in drift speed). This did not elicit prominent evoked activity and thus prohibited the reliable estimation of evoked activity. In contrast, we used a pattern reversal as stimulus change, precisely because this is known to elicit prominent evoked activity (Nakamura et al., 1997; Di Russo et al., 2005; Barnikol et al., 2006; Perfetti et al., 2007). Because of this, we were able to reliably estimate the amplitude of early visual components on single trials and demonstrate a positive correlation between gamma power and ERF amplitude, and a negative correlation between ERF amplitude and reaction times, in support of our hypothesis.

One possible concern that might confound a mechanistic interpretation of the relation between gamma-band activity, response speed, and the event-related transients could be a latent variable that correlates with these measures, causing spurious, indirect correlations. Specifically, the time interval between a warning cue and an upcoming stimulus is well known to be a determinant of response speed (Schoffelen et al., 2005; Beck et al., 2014), and temporally better predictable stimuli are associated with higher amplitudes in early components of evoked activity (Doherty, 2005; Lange et al., 2006; Dassanayake et al., 2016). Additionally, the hazard rate has been shown to correlate with spectral characteristics in alpha/beta and gamma band, (Schoffelen et al., 2005; Rolke and Hofmann, 2007; Tsunoda and Kakei, 2008; Rohenkohl and Nobre, 2011) and low frequency spectral responses have been shown to be anti-correlated with high frequency responses (Hoogenboom et al., 2006; Womelsdorf et al., 2006; Scheeringa et al., 2011; Spaak et al., 2012) Therefore, variability in the low frequency response, and/or stimulus expectancy might be a common determinant for gamma power, and response speed. We checked for these possibilities by estimating the partial correlation between gamma power and reaction time, controlling for differences in stimulus expectancy and power in low frequencies. The partial correlations were still significant, and specifically there was no additional effect of low frequency power on reaction times. This further corroborates the absence of a trial-by-trial effect of low frequency alpha/beta activity on the ERF amplitude. These results highlight the relevance and uniqueness of gamma power in behavior, and are in support of a model in which gamma-band activity modulates neuronal processing in order to affect behavior.

Even though to our knowledge there is no further literature supporting a correlation between reaction times and the amplitude of early visual evoked components, correlations have been found with their peak latency (Kammer et al., 1999; Gerson et al., 2005). In contrast to the current experiment, where the stimulus was constant in every trial and we made use of the natural variations in the physiological and behavioral response, these studies manipulated either luminance or natural scenes in order to do so. Disregarding the source of variation in the physiological and the behavioral response, it is conceivable that pre-stimulus change gamma-band activity might also affect the latency of the evoked response in addition to its amplitude, and the combination of both ultimately affects behavior. This is beyond the scope of the current study, but would be an interesting topic in future research.

Our main effect is in contrast to Privman et al. (2011). The authors used a repetition suppression paradigm, and found a reduction in ERP power in higher order visual areas as a function of gamma power in response to the second stimulus. The authors hypothesized that the gamma-band activity caused by the first stimulus might be sustained after its offset and disrupts synchronization of the neural population, selective for the second incoming stimulus. Thus, their findings might be specific to the simulation protocol used, which is further supported by the finding that the repetition suppression effect is largest when the stimuli are more similar, leading to larger overlapping neuronal representations (Grill-Spector et al., 2006).

In this study, we used non-invasive MEG recordings in human participants. In contrast to invasive recordings, MEG lacks the high spatial resolution and high signal-to-noise ratio to allow for a detailed functional and spatial interpretation of our findings. In contrast to the present findings, recent work using invasive data from macaques and cats (Ni et al., 2016), showed that the gain of the multiunit response in primary visual cortex is dependent on the gamma phase of the local field potential.

However, the authors did not investigate the functional relevance of gamma amplitude, nor did we study gamma phase. Still, their results and our results are not contradictive: the amount of synchronization on the one hand, reflected by gamma power, and high excitability phases on the other hand, might both contribute to enhanced neuronal gain.

In addition to the relatively limited spatial resolution, the high spatiotemporal variability in the response across subjects did not allow for a consistent assignment of even the early ERF components to a specific subregion in the visual system. The amplitude of the ERF was estimated as the most prominent peak within the first 100 ms after the go cue, which in terms of latency is well beyond the first geniculate input into primary visual cortex and might even reflect extrastriate activity, and thus likely reflects a more widespread activation of several cortical areas. Despite this limitation, our findings indicate that gamma-band activity increases the neuronal gain to new visual input. In addition, the fact that this effect can be shown at the spatial scale at which MEG operates, provides further justification to use gamma-band responses as a physiologically and mechanistically inspired dependent variable in non-invasive human cognitive neuroscience experiments.

## Acknowledgements

This work was supported by The Netherlands Organisation for Scientific Research (NWO Vidi grant to J.M. Schoffelen).

